# UT-018 Protects Collagen Extracellular Matrix Through Substrate-Directed Stabilization and Collagenase Modulation

**DOI:** 10.64898/2026.06.04.730073

**Authors:** Shreya Shahapur, Sadiya Mehboob, Priyanka Jadhav, Tanisha Samal, Gopi Kadiyala, Markendeya Gorantla, Uday Saxena

**Affiliations:** CDFD Technology Incubator, Centre for DNA Fingerprinting and Diagnostics (CDFD) Campus, Inner Ring Road, Uppal, Hyderabad – 500039, Telangana, India

**Keywords:** UT-018, collagenase, collagen, extracellular matrix, wound healing, matrix stabilization, regenerative medicine, dermal aging, oral care

## Abstract

Pathological collagen degradation is a central feature of impaired wound healing, dermal aging, periodontal breakdown, intestinal barrier injury and connective tissue degeneration. Current strategies often focus on direct inhibition of matrix metalloproteinases or collagenases; however, complete blockade of collagen remodeling may interfere with normal repair. UT-018, a bioactive formulation that acts as a tissue-protective and regenerative agent, was evaluated as a collagenous extracellular matrix modulator. Across in vitro kinetic assays, endpoint signal analysis, integrated area-under-curve (AUC) analysis and substrate preincubation studies, UT-018 produced concentration-dependent preservation of collagen against collagenase challenge. Importantly, collagen protection persisted after substrate preincubation with UT-018, with approximately 33%, 60% and 65% protection at 5, 10 and 25 mM UT-018 concentrations, respectively. Exploratory kinetic transformations did not support a simple competitive collagenase inhibitor model. Instead, the collective pattern supports a substrate-directed mechanism involving collagen shielding, reduced cleavage susceptibility and indirect modulation of collagenase activity. These findings position UT-018 as a potential first-in-class collagen resilience modulator for wound healing, gastrointestinal barrier protection, oral care, dermal preservation and regenerative medicine applications.

**Highlights:**

- UT-018 preserves collagen content in in vitro collagenase challenge assays.
- The Protection is UT-018 concentration-dependent across kinetic, endpoint and AUC readouts.
- Preincubation of substrate with UT-018 retains protection after collagenase challenge.
- The data support matrix-directed stabilization by UT-018 rather than classical active-site collagenase inhibition.

## Introduction

Collagen is the major structural protein of the extracellular matrix (ECM) and provides tensile strength, tissue architecture and a scaffold for repair. Physiological collagen remodeling is essential for wound healing, angiogenesis and tissue renewal. However, excessive collagenase and matrix metalloproteinase activity can destabilize the ECM and contribute to chronic wounds, diabetic ulcers, periodontal disease, inflammatory bowel disease, skin aging, osteoarthritis and connective tissue degeneration.

Most anti-collagenolytic approaches have focused on enzyme inhibition. While this strategy can reduce degradation, broad suppression of collagenases may be biologically undesirable because controlled collagen turnover is required for normal remodeling. A more attractive strategy is to increase collagen resilience to pathological degradation without fully shutting down protease activity.

UT-018 is proposed as a regenerative and tissue-stabilizing platform. The central hypothesis tested here is that UT-018 protects collagenous substrates through matrix-directed stabilization and substrate shielding rather than through a classical active-site collagenase inhibition mechanism.

## Materials and Methods

Collagenase activity study design. UT-018 was evaluated in in vitro collagenase-mediated degradation assays using commercial assay kits. Parameters examined were kinetic, endpoint, integrated AUC and substrate preincubation outcomes. The analysis was designed to determine whether UT-018 preserves collagen signal in a concentration-dependent manner and whether protection persists when the substrate is exposed to UT-018 before collagenase challenge.

Kinetic collagenase assay. Kinetic data were generated in replicate studies. Blank-corrected signal was monitored over time and summarized as mean kinetic traces with variation bands. Higher retained signal was interpreted as greater preservation of collagenous substrate.

Endpoint signal analysis. Final blank-corrected signal was compared across conditions. Pairwise comparisons were used to assess whether increasing UT-018 exposure produced statistically distinguishable collagen preservation.

Integrated protection analysis. Area under the kinetic trace was calculated to capture the total retained collagen signal over the full assay period. This readout was used to distinguish sustained protection from transient endpoint effects.

Mathematical kinetic interpretation. Lineweaver-Burk and Michaelis-Menten style visualizations were used as exploratory tools to compare the observed profile with classical competitive, noncompetitive and substrate-directed stabilization models.

Substrate preincubation assay. Collagen or gelatin substrate was preincubated with UT-018 before collagenase challenge. Persistence of protection after preincubation was used as support for substrate-directed stabilization. A protection signal that remains after substrate exposure is more consistent with collagen or matrix stabilization than with transient enzyme active-site blockade.

## Results

### UT-018 preserves collagen signal across the kinetic collagenase assay

Mean kinetic traces showed (Figure 1) a clear separation among the conditions over the full assay time frame. The lowest condition (lowest concentration of UT-018, 5 mM) maintained the lowest blank-corrected collagen signal, while the intermediate and highest conditions showed progressively greater preservation. The highest condition maintained the greatest OD signal throughout the time course, indicating that UT-018 protection was not a late endpoint artifact but was present across the kinetic phase of collagenase exposure. The relatively stable separation among UT-018 conditions supports sustained matrix protection.

**Figure 1.**
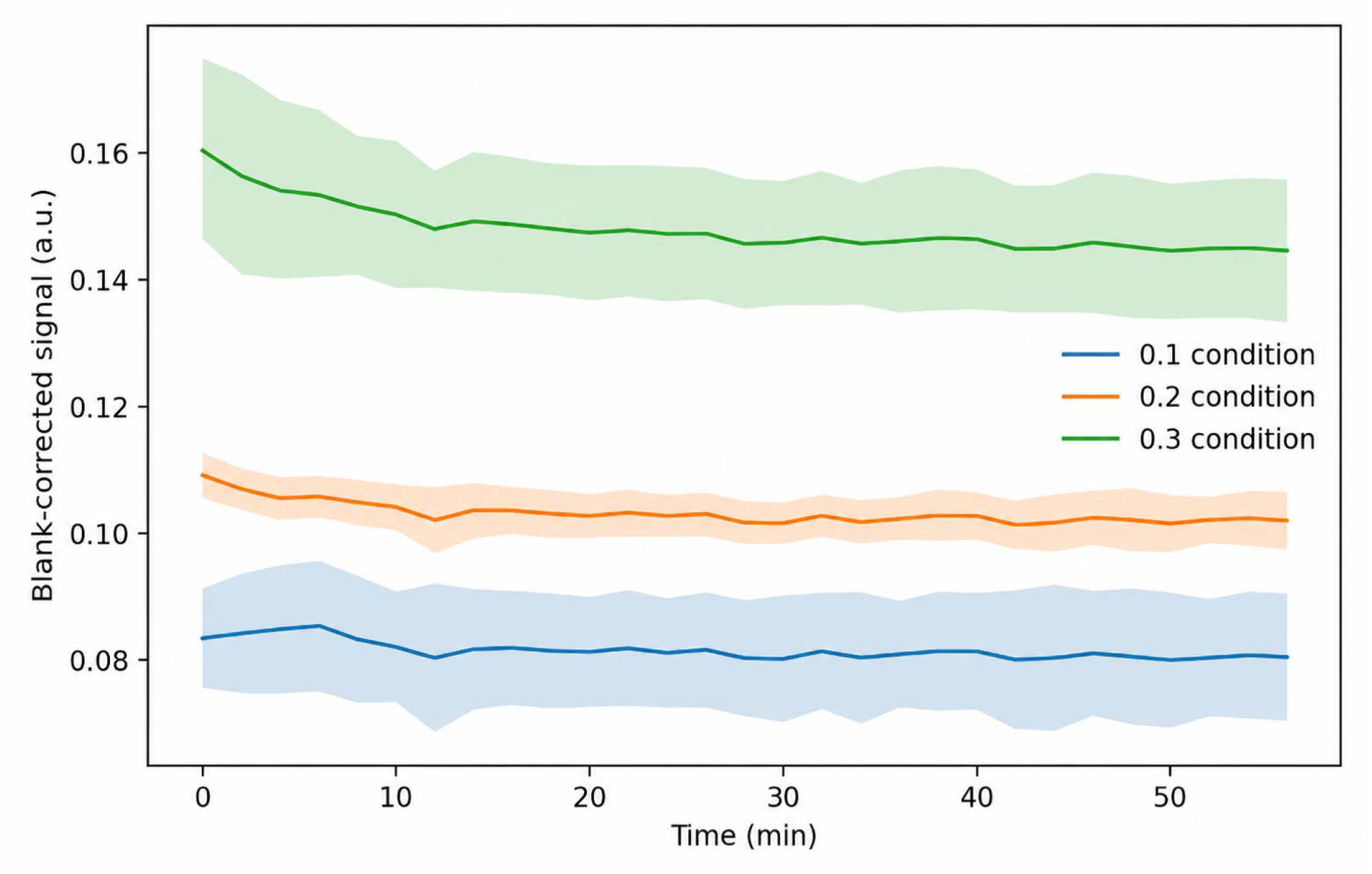
Mean kinetic traces demonstrating concentration-dependent collagen preservation by UT-018 (5, 10 and 25 mM, shown above as 0.1, 0.2, and 0.3 conditions). Blank-corrected collagen signal was followed over time using replicate collagen sheets. The highest concentration of UT-018 tested showed the greatest retained signal across the assay window, while the intermediate condition showed partial preservation. This kinetic pattern supports sustained collagen protection rather than a transient endpoint effect.

### Endpoint analysis confirms graded collagen protection

Final endpoint signals increased (Figure 2) in a graded fashion across concentrations of UT-018, rising from approximately 0.081 at the lowest condition to approximately 0.102 at the intermediate condition and approximately 0.145 at the highest condition. Pairwise comparisons demonstrated statistically significant separation between the lowest and intermediate conditions, the intermediate and highest conditions, and the lowest and highest conditions. This confirms that UT-018 produced concentration-dependent preservation of collagen signal at the assay endpoint.

**Figure 2.**
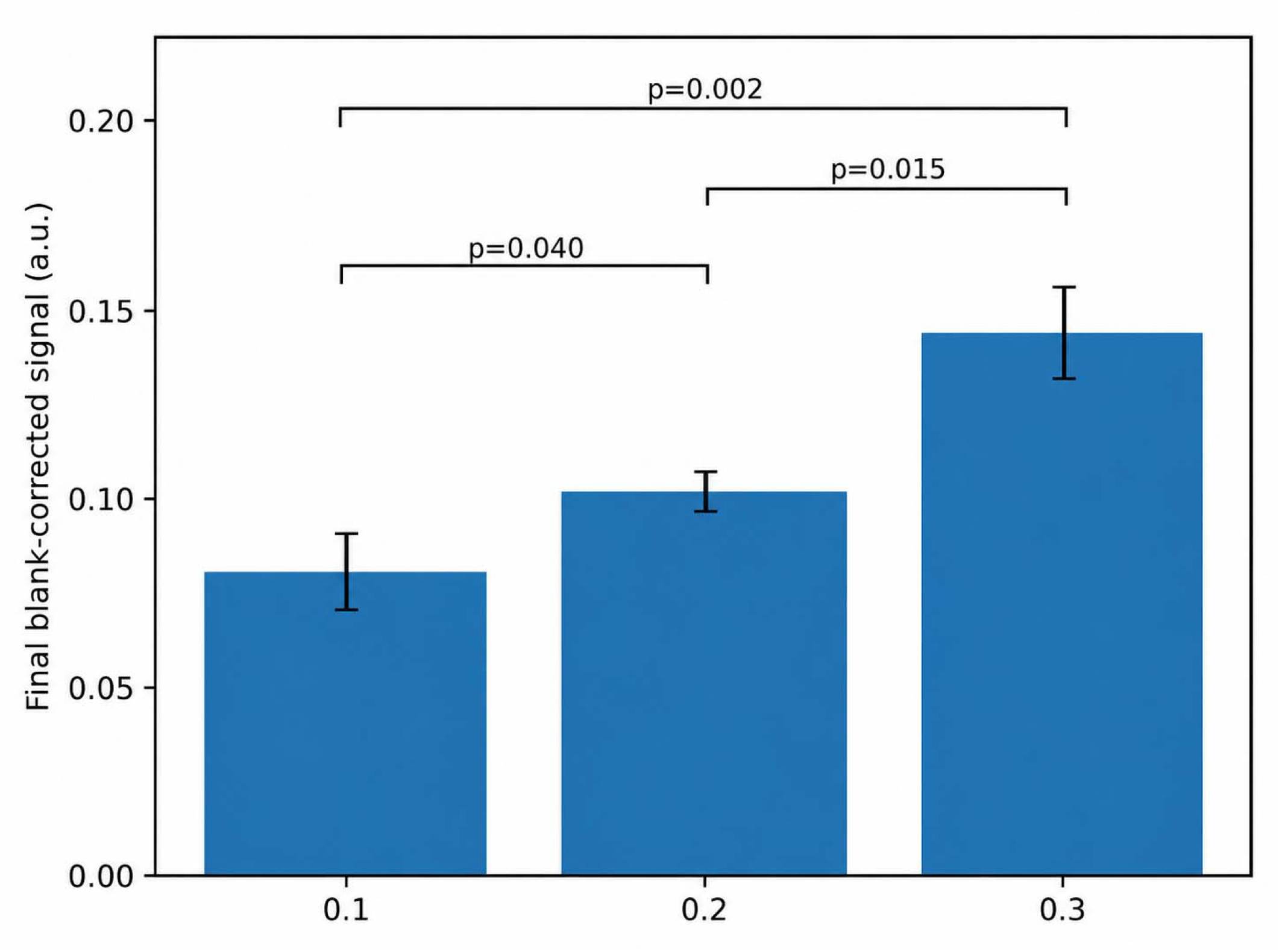
Endpoint collagen preservation following exposure to UT-018 (5, 10 and 25 mM, shown above as 0.1, 0.2, and 0.3 conditions). Final blank-corrected signal increased across the workbook-listed conditions. Reported pairwise comparisons were p = 0.040 for low versus intermediate, p = 0.015 for intermediate versus high, and p = 0.002 for low versus high. These data support a graded and statistically significant collagen preservation effect by UT-018.

## Integrated AUC analysis demonstrates sustained protection over time

Because a single endpoint can miss time-dependent differences, integrated AUC was used to quantify total retained signal over the full assay. As shown in Figure 3, AUC increased from approximately 4.72 to 5.95 and 8.55 across increasing conditions. The high concentration condition was significantly greater than both lower conditions, indicating a sustained protection effect throughout the assay. The intermediate condition trended above the lowest condition, consistent with partial protection.

**Figure 3.**
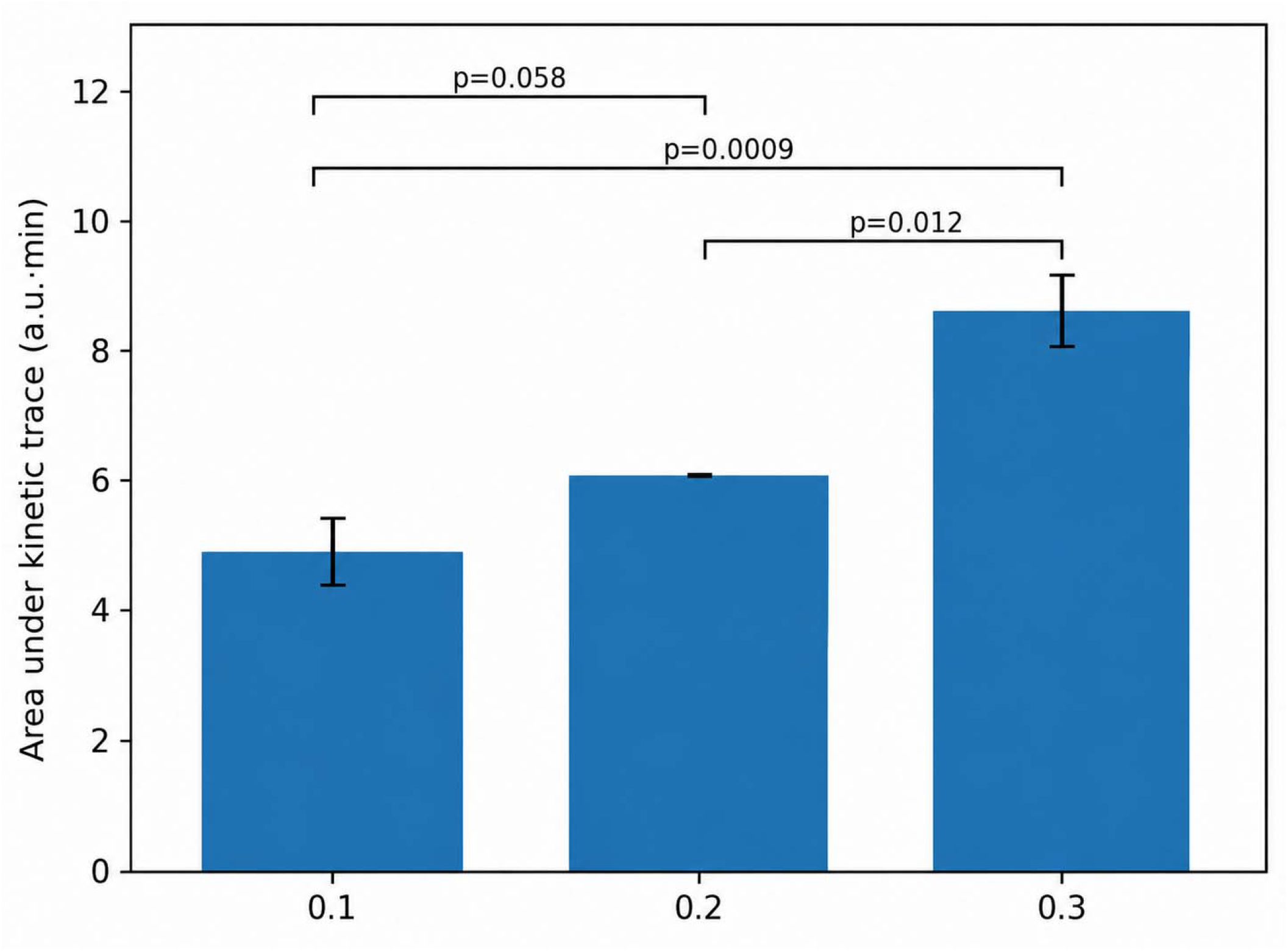
Integrated AUC analysis of collagen protection by UT-018 (5, 10 and 25 mM, shown above as 0.1, 0.2 and 0.3 conditions). AUC increased across conditions, indicating that UT-018 preserved collagen signal over the full assay period. Reported comparisons included p = 0.058 for low versus intermediate, p = 0.012 for intermediate versus high, and p = 0.0009 for low versus high. This analysis strengthens the interpretation that UT-018 provides sustained matrix protection.

### Exploratory Lineweaver-Burk transformation does not support simple competitive inhibition

An apparent Lineweaver-Burk transformation was used as an exploratory mechanistic view. The pattern (Figure 4) did not cleanly match a classical competitive inhibitor profile. In a simple competitive model, one would expect a predictable shift in apparent Km with a relatively preserved Vmax. The collagen substrate system with UT-018 instead showed a profile more consistent with altered substrate susceptibility and matrix-protease interaction rather than exclusive active-site competition.

**Figure 4.**
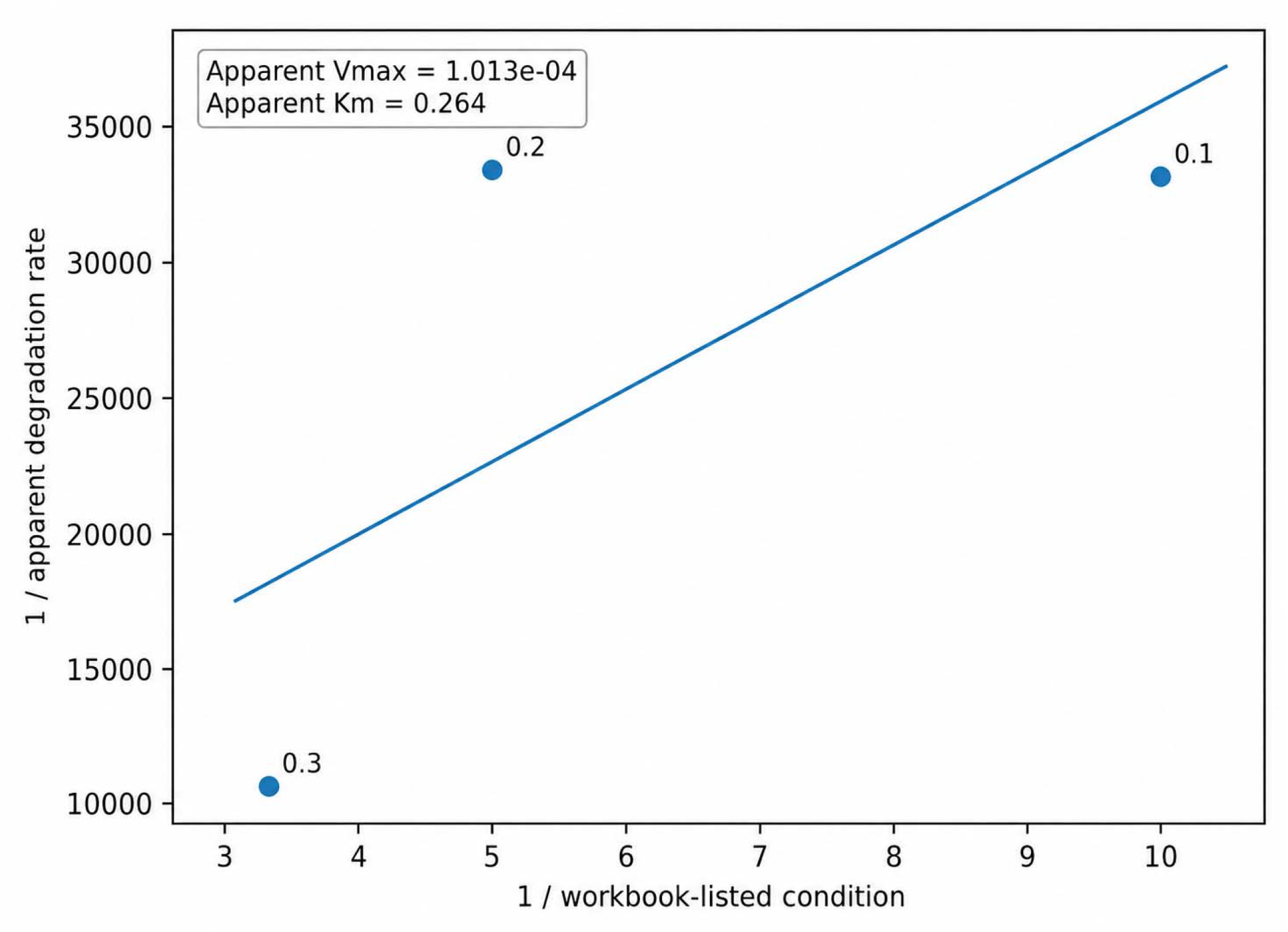
Exploratory Lineweaver–Burk transformation of collagenase assays performed with UT-018 (5, 10 and 25 mM, shown above as 0.1, 0.2 and 0.3 conditions). The apparent kinetic profile is not consistent with a simple classical competitive inhibitor model. Because the assay uses collagenous gel substrate rather than a purified soluble substrate, this plot should be interpreted as a qualitative mechanistic aid. The pattern supports matrix-directed protection and altered cleavage susceptibility.

### Michaelis-Menten model interpretation favors substrate-directed stabilization

A conceptual Michaelis-Menten interpretation was used to distinguish direct enzyme inhibition from matrix stabilization. Classical competitive inhibition, noncompetitive inhibition and substrate stabilization generate different curve shapes. Shown in Figure 5, the UT-018 dataset is most consistent with a model in which collagen becomes less susceptible to enzymatic cleavage without complete inhibition of the collagenase system. This interpretation is biologically important because partial preservation of ECM integrity may allow physiological remodeling to continue.

**Figure 5.**
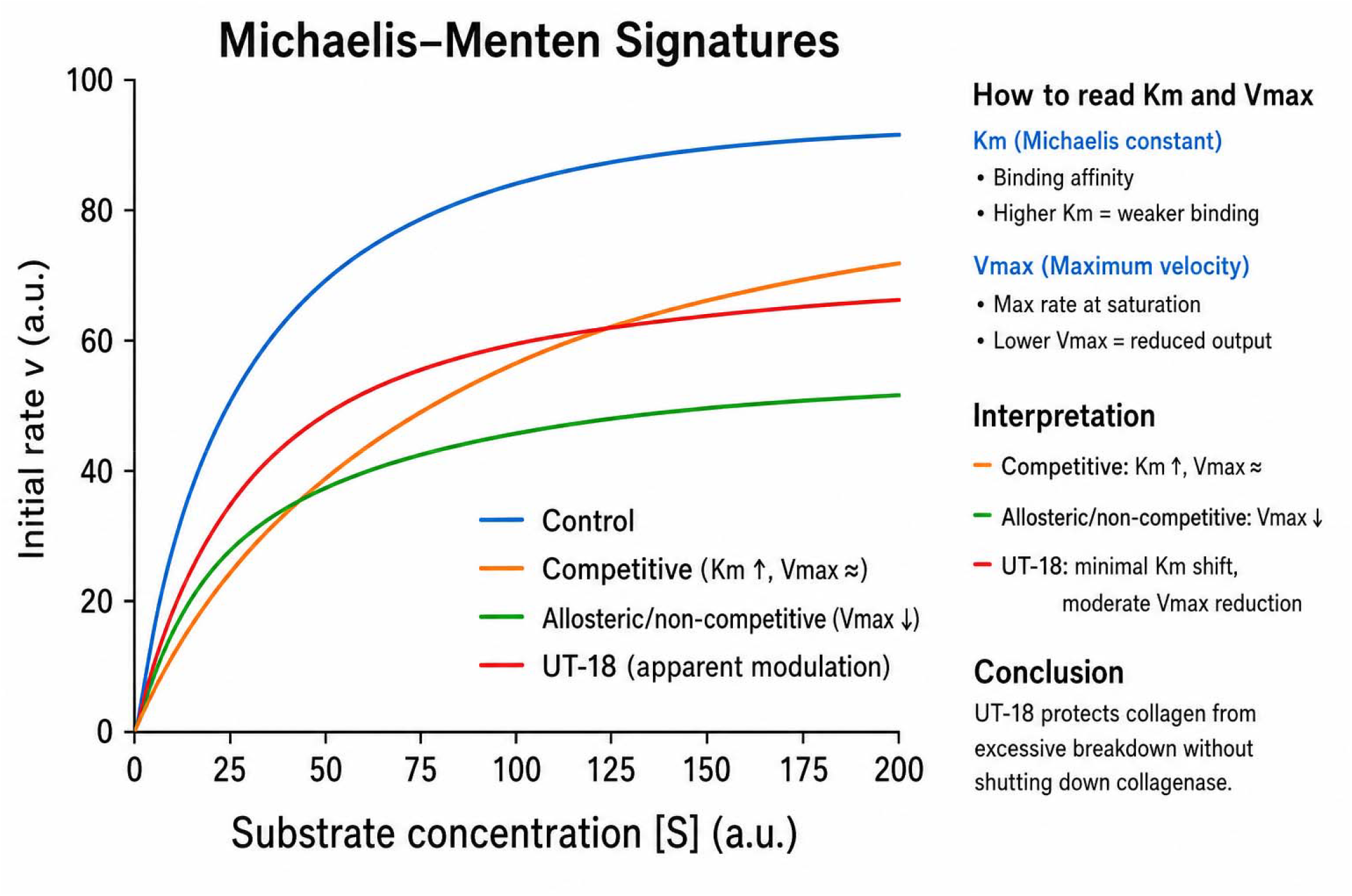
Michaelis–Menten interpretation of collagen stabilization by UT-018 (5–25 mM). The interpretive model compares classical competitive inhibition, noncompetitive inhibition and substrate stabilization. The UT-018 profile aligns best with collagen or matrix stabilization, where the substrate becomes less accessible or less vulnerable to collagenase-mediated cleavage while enzyme throughput is not fully abolished.

### Substrate preincubation demonstrates persistent collagen protection

We next tested whether reduced collagen degradation would still be observed after preincubating the collagenous substrate with UT-018 prior to collagenase exposure. The substrate preincubation assay provides the strongest mechanistic evidence for a matrix-directed effect. UT-018 produced approximately 33% protection at 5 mM, approximately 60% protection at 10 mM and approximately 65% protection at 25 mM after preincubation (Figure 6). The response approached a plateau between 10 and 25 mM, suggesting a saturable interaction with collagenous substrate or matrix microdomains. Persistence of protection following preincubation is difficult to explain by simple transient enzyme active-site inhibition and instead supports potentially substrate shielding, altered hydration, ionic stabilization or reduced collagenase accessibility.

**Figure 6.**
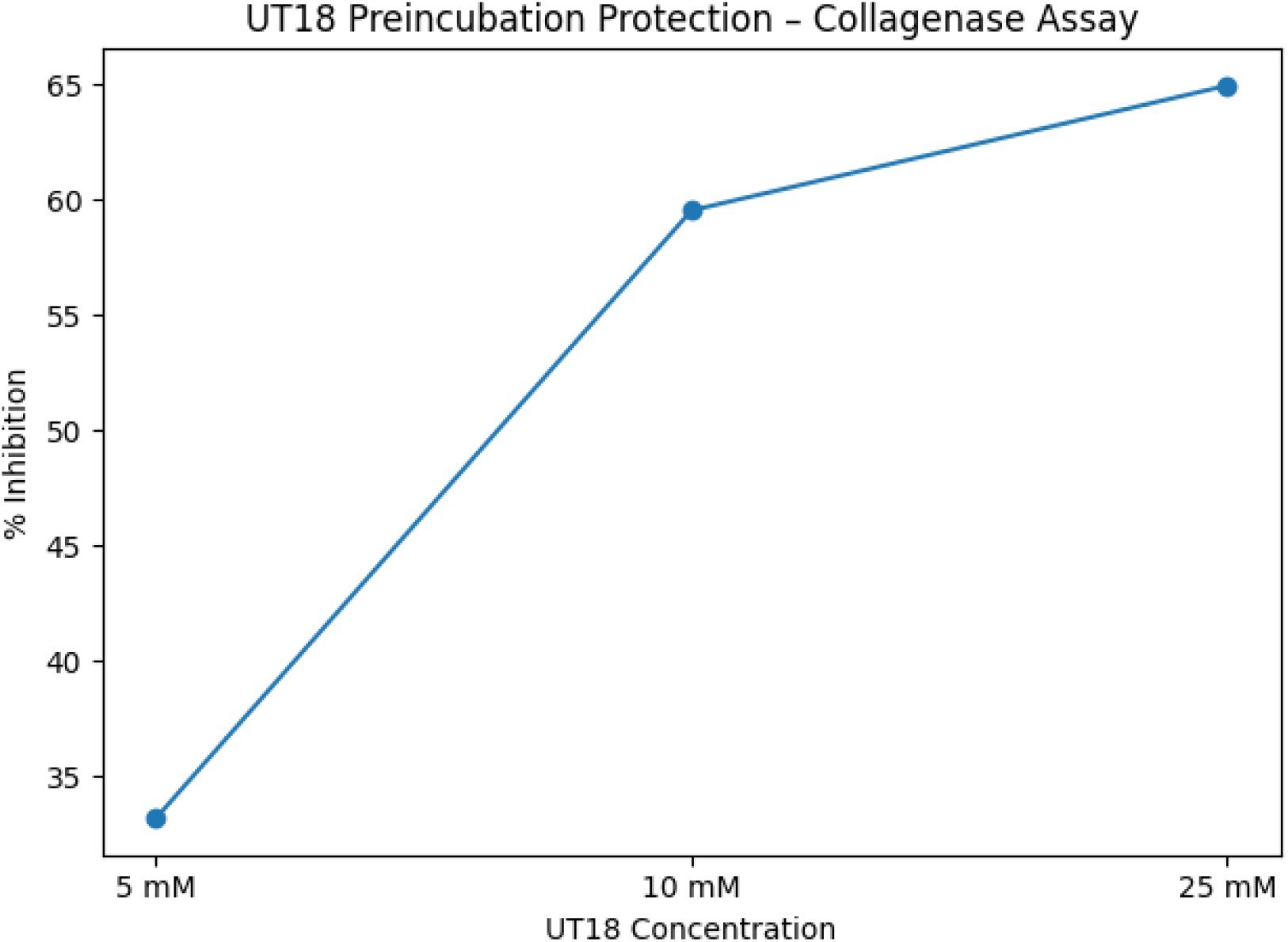
Substrate preincubation protection assay demonstrating matrix stabilization by UT-018 at 5, 10 and 25 mM. UT-018 protected collagenous substrate after preincubation, with approximately 33%, 60%, and 65% protection against collagen degradation at 5, 10 and 25 mM, respectively. The plateau between 10 and 25 mM suggests saturable matrix stabilization. This is the key finding supporting a substrate-directed mechanism.

### Integrated mechanistic interpretation

Taken together, the kinetic, endpoint, AUC and preincubation studies support a coherent mechanism: UT-018 increases collagen resilience to enzymatic cleavage. The findings are more consistent with extracellular matrix stabilization than with direct, complete collagenase blockade.

| Observation | Interpretation | Relevance |
| --- | --- | --- |
| Separated kinetic traces | Protection occurs throughout the time course | Not only an endpoint artifact |
| Endpoint signal increases | Concentration-dependent preservation | Supports graded UT-018 activity |
| AUC increases | Sustained integrated protection | Confirms durability over assay window |
| Lineweaver-Burk profile not classical | Not simple competitive inhibition | Suggests noncanonical mechanism |
| Preincubation protection persists | Substrate-directed matrix stabilization | Strongest mechanistic support |

## Discussion

Our data support UT-018 as a collagenous ECM resilience modulator rather than a direct inhibitor of the collagenase enzyme. The main finding is that UT-018 preserves collagen signal across kinetic, endpoint and integrated AUC readouts and that this protection persists after substrate preincubation. The preincubation result is particularly important because it indicates that UT-018 can protect the collagenous substrate itslf.

The mechanism is best described as substrate-directed stabilization rather than classical collagenase inhibition. This distinction matters translationally. A direct collagenase inhibitor may suppress pathological degradation but may also impair normal remodeling. A matrix stabilizer could instead preserve tissue architecture during excessive protease exposure while still allowing physiological collagen turnover.

The observed plateau in the preincubation assay suggests that UT-018 may interact with finite collagen binding or matrix hydration sites. This feature could reduce collagenase access or cleavage susceptibility without requiring complete enzyme inhibition.

### Translational significance

The collagen-preserving effects of UT-018 observed in the present study provide a mechanistic framework that may partly explain the regenerative outcomes previously reported in independent wound-healing and hair-regeneration models (Saxena et al., 2026). In those studies, topical UT-018 application accelerated wound closure, enhanced granulation tissue formation, improved collagen organization, increased fibroblast proliferation, promoted angiogenesis, and stimulated follicular regeneration with increased follicular density and anagen-phase transition. Histological analyses consistently demonstrated preserved extracellular matrix architecture and organized stromal remodeling following UT-018 treatment. The present findings suggest that direct stabilization of collagenous substrates and reduced susceptibility to collagenase-mediated degradation may contribute at least in part to these in vivo effects by preserving the structural scaffold required for cellular migration, tissue repair, angiogenesis, and follicular remodeling.

The UT-018 collagenase dataset shown here supports potential applications in several tissue repair settings. In chronic wounds and diabetic ulcers, excessive protease activity degrades newly deposited matrix and delays closure. In oral care, collagen breakdown contributes to gingival recession and periodontal tissue loss. In dermal aging, collagen fragmentation reduces elasticity and structural integrity. In gastrointestinal disorders, preservation of ECM and barrier-associated matrix may support mucosal resilience. These applications should be further developed with indication-specific assays, including wound fluid protease models, human dermal matrix systems, gingival fibroblast matrix models and intestinal barrier injury models.

## Conclusion

UT-018 demonstrates concentration-dependent protection of collagenous substrates from collagenase-mediated degradation. The strongest mechanistic evidence comes from substrate preincubation, where protection persisted and approached saturation. The collective data support UT-018 as a matrix-directed collagen resilience modulator with potential relevance to wound healing, dermal preservation, oral care, gastrointestinal barrier protection and regenerative medicine.

